# Host adaptation mediated by intergenic evolution in a bacterial pathogen

**DOI:** 10.1101/236000

**Authors:** S. M. Hossein Khademi, Lars Jelsbak

**Affiliations:** Department of Biotechnology and Biomedicine, Technical University of Denmark, 2800 Lyngby, Denmark

**Keywords:** intergenic evolution, *cis*-regulatory elements, bacterial adaptation, gene expression, antibiotic resistance

## Abstract

Bacterial pathogens evolve during the course of infection as they adapt to the selective pressures that confront them inside the host. Identification of adaptive mutations and their contributions to pathogen fitness remain a central challenge. Although mutations can either target intergenic or coding regions in the pathogen genome, studies of host adaptation have focused predominantly on molecular evolution within coding regions, whereas the role of intergenic mutations remains unclear. Here, we address this issue and investigate the extent to which intergenic mutations contribute to the evolutionary response of pathogens to host environments, and whether intergenic mutations have distinct roles in host adaptation. We characterize intergenic evolution in 44 lineages of a clinically important bacterial pathogen, *Pseudomonas aeruginosa*, as they adapt to their hosts. We identify 88 intergenic regions in which parallel evolution occurs. At the genetic level, we find that mutations in these regions under selection are located primarily within regulatory elements upstream of transcriptional start sites. At the functional level, we show that these mutations both create or destroy regulatory interactions in connection to transcriptional processes and are directly responsible for evolution of important pathogenic phenotypes including antibiotic sensitivity. Importantly, we find that intergenic mutations are more likely to be selected than coding region mutations and that intergenic mutations enable essential genes to become targets of evolution. In summary, our results highlight the evolutionary significance of intergenic mutations in creating host-adapted variants and that intergenic and coding regions have different qualitative and quantitative contributions to this process.

**Significance:** Pathogens adapt to their host during infection, but the contribution and function of non-coding intergenic sequences to adaptation is poorly understood. Here, genome-wide identification of adaptive mutations within intergenic regions demonstrates that these sequences constitute an important part of the genetic basis for host adaptation. We find that intergenic mutations are abundant relative to adaptive mutations within coding sequences, and can contribute directly to evolution of pathogen-relevant traits. Importantly, we find that intergenic mutations modify expression of essential genes and thus make contributions that are functionally distinct from coding mutations. These results improve our understanding of the evolutionary processes *in vivo* and can potentially assist in refining predictions of pathogen evolution, disease outcome and antibiotic resistance development.

## Introduction

Bacterial pathogens evolve during infection as they adapt to the environment inside the host (1). Since the bacterial phenotypes selected *in vivo* may have profound impact on disease severity and progression (2, 3) as well as response to antibiotic therapy (4), identification and analysis of the full range of beneficial genetic changes that underlies host adaptation is of importance.

Although adaptive mutations may potentially change the sequences of either coding regions or non-translated intergenic regions and thus affect protein function or expression, respectively, studies of pathogen adaptation during infection of host tissues have focused predominantly on molecular evolution within coding regions, whereas the role of adaptive mutations in intergenic regions has received comparably less attention. The shortage of systematic, genome-wide analyses of intergenic evolution in bacterial pathogens is surprising, given the fact that these regions are home to a large number of functional elements required for expression of virulence and resistance determinants *in vivo* and that intergenic regions are maintained by purifying selection in many bacterial species (5–7). Moreover, *cis*-regulatory mutations are known to play an important role in phenotypic evolution in eukaryotic organisms (8). Overall, it remains unclear to what extent intergenic mutations contribute to the evolutionary response of pathogens to the host environment and whether intergenic mutations have a qualitative distinct role in host adaptation.

There are clear, albeit few, examples of intergenic regions that evolve under selection within the host. For example, evolution of a novel regulatory interaction between the virulence regulator SsrB and the promoter of the *sfrN* gene was shown to result in enhanced within-host fitness in *Salmonella* Typhimurium (9). In *Pseudomonas aeruginosa*, evolution of the intergenic region of the *phuR-phuSTUVW* genes during host colonization was shown to increase expression of the Phu heme uptake system and improve the ability of the pathogen to acquire iron from hemoglobin (10). In *Mycobacterium tuberculosis*, evolution of ethambutol resistance has been linked to the *empAB* promoter region acquiring mutations that enhance expression of enzymes essential for the synthesis of cell wall arabinogalactan (11). These and other examples point towards an evolutionary significant role of mutations in intergenic regions in connection to bacterial pathogenesis and justify a broader analysis of this type of mutations.

One reason for the paucity in genome-wide analysis of intergenic evolution is probably related to the inherent difficulties in inferring function directly from the sequence within intergenic regions and, consequently, to differentiate adaptive mutations with functional effects from neutral mutations that have been fixed by chance. Here, we harnessed the combination of parallel evolution and functional genomics to identify intergenic regions under selection in the genome of the opportunistic pathogen *P. aeruginosa* during the process of host adaptation in multiple cystic fibrosis patients. Our study reveals that adaptive intergenic mutations represent an egregiously underappreciated aspect of host adaptation in *P. aeruginosa*, and that intergenic and coding region mutations contribute differently to this process both qualitatively and quantitatively.

## Results

### Parallel evolution in intergenic regions in P. aeruginosa

To investigate the contribution of intergenic mutations to bacterial adaptation to the host environment, we considered data from seven studies (12–18) in which multiple clonal *P. aeruginosa* isolates have been sampled and sequenced during the course of infection in subjects with cystic fibrosis (CF). In CF infections, the host environments in individual subjects represent parallel selective conditions by which evolution is directed, and identification of parallel genetic evolution in bacteria from independent infections is strongly suggestive of positive selection at these loci (16).

Here, we focused our analysis exclusively on intergenic regions in which mutations were acquired during infection and included only intergenic regions also present in the PAO1 reference genome. In total, we identified 3,489 mutations (2,024 SNPs and 1,465 indels) in the intergenic regions of the 44 different *P. aeruginosa* clone types included in our dataset (Table S1). Since the majority of regulatory elements in the bacterial genome range between 5-30 bp in length (19), we identified intergenic regions under positive selection by only considering mutations found in parallel in different clone types and distributed within a window of less than 30 bp (Materials and Methods). Applying these criteria, we found 63 intergenic regions with parallel genetic evolution (Figure 1).

**Figure 1:**
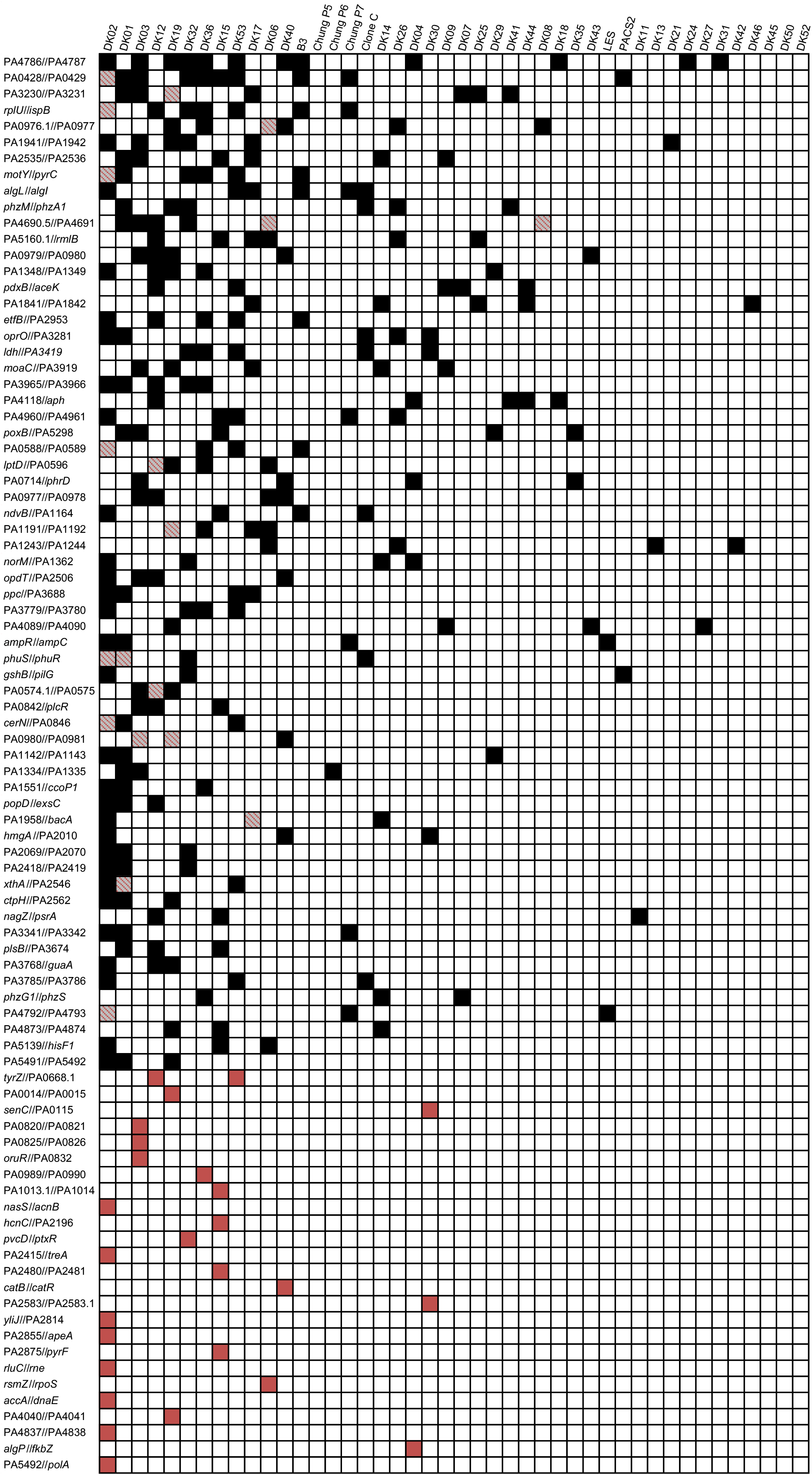
Pathoadaptive intergenic regions. Regions targeted by mutations involved in host adaptation through parallel evolution across or within clone types. The black squares in the matrix show the intergenic region with parallel mutations in isolates of the respective clone type. The red squares in the matrix show the intergenic region with parallel mutations within isolates of a clone type alone. Squares with striped red color indicate regions in which mutations has been selected both within isolates of that clone type and across other clone types.

Since certain *P. aeruginosa* clone types are transmissible and can form clinic-specific outbreaks among patients (16), we also analyzed if distinct intergenic mutations had accumulated in parallel among clonal isolates within each of the 44 clone types. We identified 41 intergenic regions in which three or more distinct mutations (less than 30 bp apart) had accumulated in isolates of the same clone type (Figure 1). Interestingly, 16 of these regions are also represented among the 63 regions identified in our analysis of parallel mutations between clone types, providing further support for the importance of these mutations in adaptation of *P. aeruginosa* to the CF environment (Figure 1). In total, we identify 88 intergenic regions that evolved under the pressure of natural selection within the hosts. The connection between these ‘pathoadaptive’ regions and their flanking genes identify genetic systems with importance for pathogen adaptation and thus provide insight into the selective forces that operate on the pathogen.

### Intergenic mutations frequently target promoter sequences

We next analyzed the genomic distribution of the identified intergenic mutations. Non-translated intergenic regions are distributed across the genome in three possible orientations: 1) upstream of two genes, 2) downstream of two genes and 3) upstream of one gene and downstream of one gene, where the latter may include regions with no promoter and within an operon (Figure 2a). We found an over-representation of mutations upstream of two genes among the pathoadaptive regions (Fisher’s exact test, *P* = 0.010, *n* = 88, Figure 2b). This bias towards selection of intergenic mutations upstream of genes suggests that the majority of intergenic mutations target potential *cis*-regulatory elements such as the core promoter, transcription factor binding sites, ribo-regulators or translational elements, and consequently influence protein expression levels by affecting transcriptional or post-transcriptional processes.

**Figure 2:**
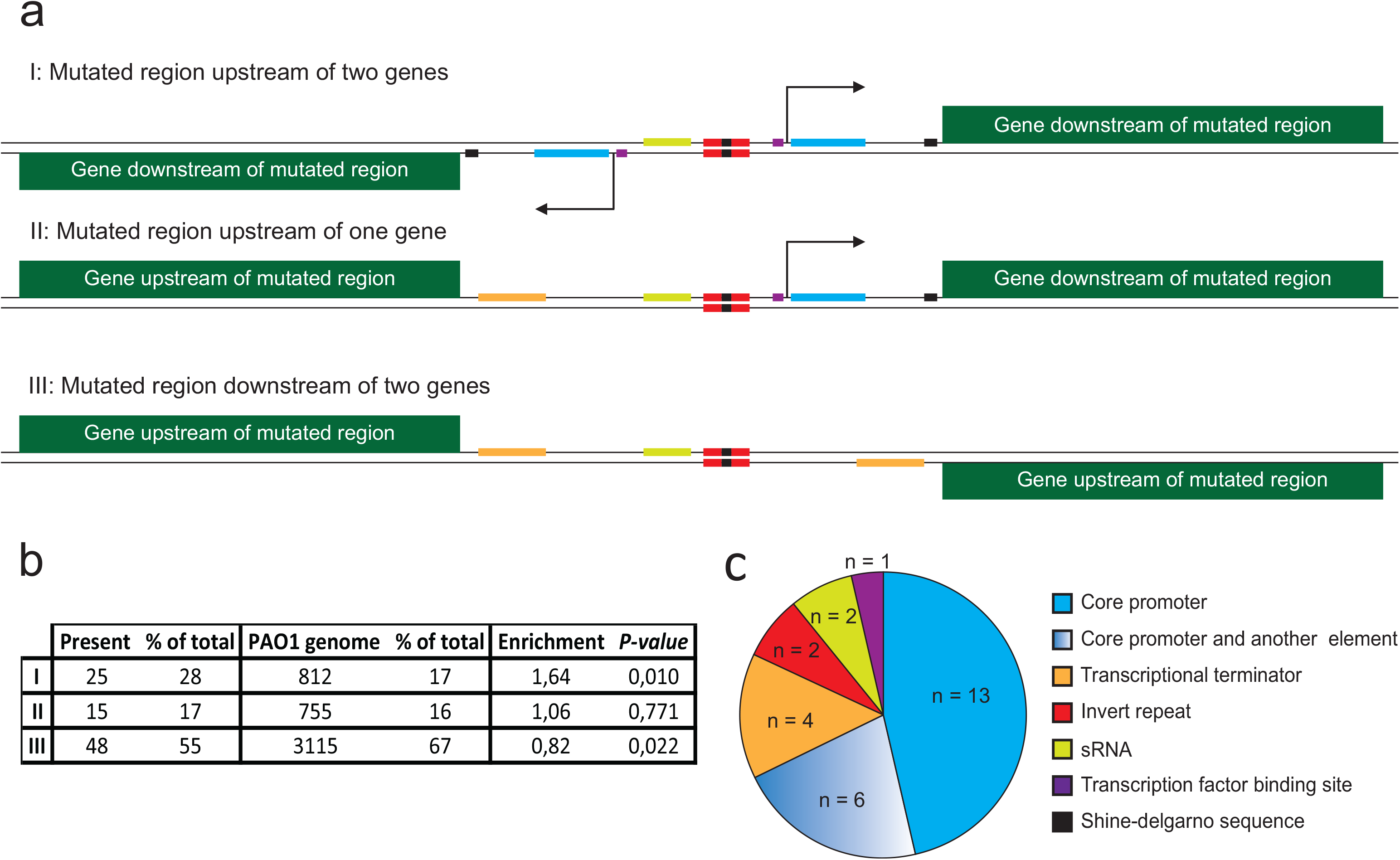
Orientation and regulatory elements in intergenic regions. a) Overview of the three different orientations of intergenic regions and the possible location of potential elements within each type. b) Distribution of different orientations of intergenic regions (I-III) within PAO1 genome and the pathoadaptive intergenic regions. Two-tailed Fisher’s exact test is performed to analyze over-representation or under-representation of certain orientations within pathoadaptive intergenic regions (n = 88). c) Pie chart demonstrating the distribution of putative intergenic elements targeted by pathoadaptive intergenic mutations among regions where the mutation cluster was within any known element (n = 28).

To further explore this hypothesis, we analyzed the complete set of 88 pathoadaptive regions for the presence of known regulatory elements (Materials and Methods) and mapped the overlap between these putative regulatory sites and the identified adaptive mutations. While bacterial intergenic regions are home to a wide range of regulatory elements many of which are not well characterized, we nevertheless observed 28 regions (32%), in which the cluster of adaptive mutations was positioned within one or more putative regulatory elements. The majority of mutations within these 28 regions target the putative core promoter alone or in combination with other elements (Figure 2c), suggesting that intergenic mutations frequently target sequences important for transcriptional processes. In support of this, we observed that intergenic mutations were more frequently located upstream of known transcriptional start sites (TSS) (37 cases) than downstream (10 cases) (Table S6).

### Pathoadaptive intergenic mutations change transcriptional activity of genes involved in host interaction, metabolism and antibiotic susceptibility

To further explore the potential relationship between intergenic mutations and transcription, we quantified the effects of a subset of intergenic mutations on transcription of downstream genes. To this end, we constructed transcriptional fusions of both wild type and mutant intergenic alleles with the luciferase reporter (*luxCDABE*) genes and integrated single copies of the fusions at the neutral *attB* site (10) in the chromosome of *P. aeruginosa* PAO1 (20). We measured the transcriptional activity of 25 different intergenic regions in which pathoadaptive mutations were located upstream of either one or two genes. This selection resulted in a total of 32 transcriptional fusions, which represent 33% of all possible fusions within the complete set of 88 pathoadaptive regions. In addition, for one of the intergenic regions (*ampR//ampC*), we tested two alleles, each with different mutations (Table S7 and Figure S1).

Measurements of *lux* expression during exponential growth in Luria-Bertani (LB) medium and ABTGC minimal medium revealed significantly altered expressions in 16 of 34 tested fusions in at least one of the two conditions (Student *t* test, *P* < 0.05, Figure 3). Altered expression was in most cases moderate (<3-fold change) and the fold change ranged from −3.1 to 22.1 for the mutant allele compared to that of wild type (Figure 3). Interestingly, 10 of these 16 fusions exhibited altered expressions only in either LB or ABTGC minimal medium, but not in both conditions, which suggests that many adaptive intergenic mutations alter transcriptional levels while not interfering with conditional control mechanisms.

**Figure 3:**
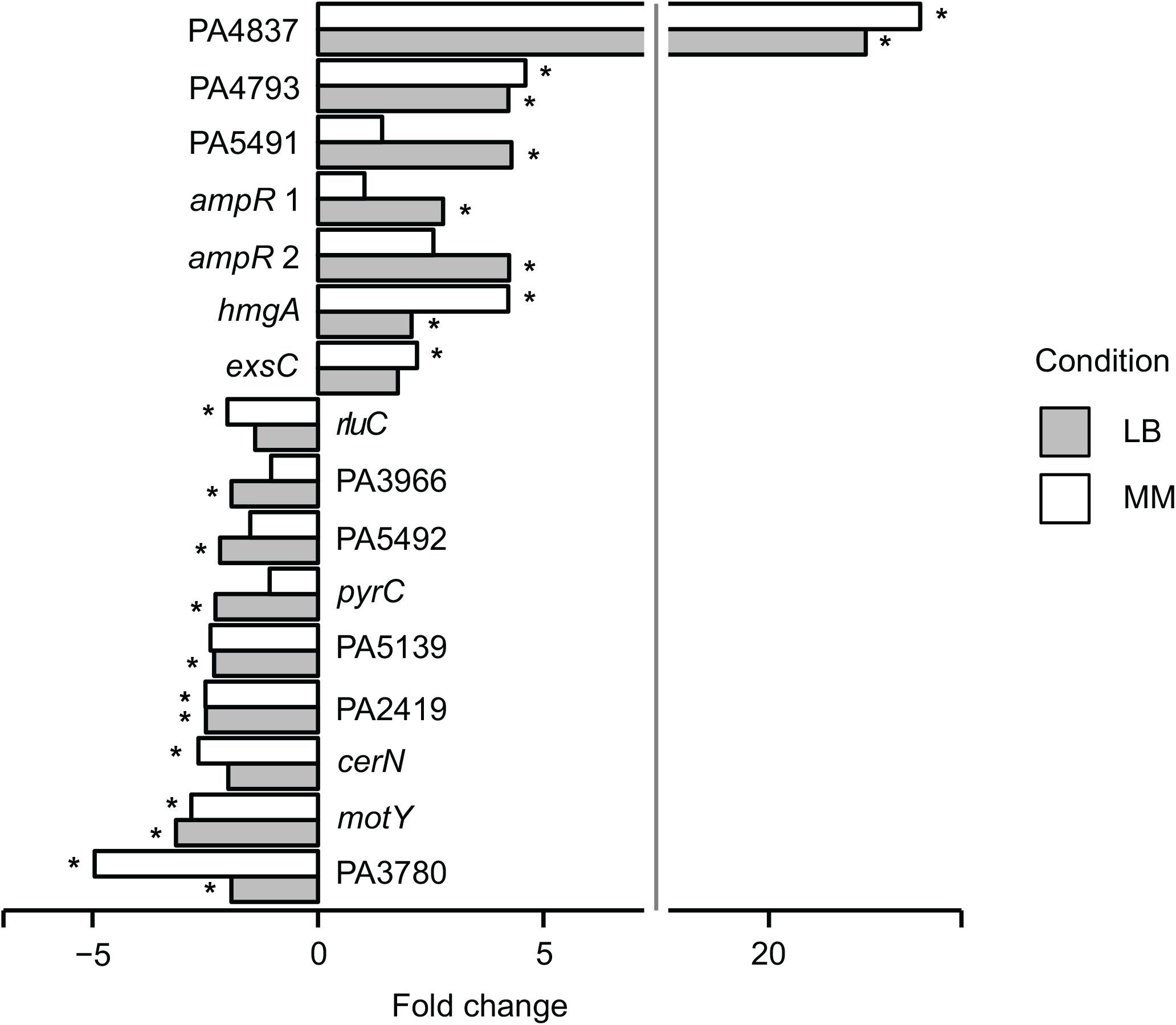
Intergenic mutations with functional effects on transcription. Expression of *lux* from transcriptional fusions with mutated and wild type alleles were measured at OD_600_ = 0.15 and normalized by cell density. Transcriptional fusions were examined in Luria-Bertani (LB) and ABTGC minimal media. Mean luminescence was calculated for three biological replicates and the relative fold change of mutant versus wild type allele calculated. Statistical analysis of the difference between two means was performed by a two-tailed Student *t* test and the asterisk denotes *P* < 0.05.

These results show that a substantial fraction of the intergenic mutations are associated with functional (transcriptional) effects despite the fact that we recorded these effects in the non-native PAO1 genetic background (*i.e*. with removal of potential epistatic effects from the additional mutations found in the clinical isolate) and in a narrow range of conditions, which most likely means that we are not capturing the full spectrum of functional effects connected to the intergenic mutations.

Several of the 16 fusions with altered expression relate to genes that encode proteins with known functions in bacteria-host interactions, cellular metabolism and antibiotic resistance. For example, *cerN* expresses a ceramidase involved in utilization of host-produced sphingolipids (21), *exsC* expresses a protein involved in positive regulation of the type III secretion system (22) and PA4837 is the first gene in an operon (PA4837-34) involved in expression of a metalophore system essential for survival in airway mucus secretions (23, 24). Other genes are known to play a role in pyrimidine and aromatic amino acid metabolism (*pyrC* and *hmgA*, respectively). Finally, two genes are linked to antibiotic resistance: *rluC* (25) and *ampR* (26). Seven genes encode proteins of unknown functions and their role in relation to host adaptation remains unclear.

Interestingly, expression changes were observed in both directions (seven mutant alleles resulted in increased expression and nine mutant alleles resulted in decreased expression) (Figure 3), suggesting that pathoadaptive intergenic mutations may equally well either create or destroy regulatory interactions.

### Mutations upstream of ampR and ampC enhance resistance to several antibiotics

Next, we explored the direct effects of intergenic mutations on the physiology of the pathogen. As resistance towards antibiotics is a common phenotype that emerges during CF infections, we selected the mutations found in the two alleles of the *ampR//ampC* intergenic region for further study. Mutations in this intergenic region resulted in enhanced expression of the global antibiotic resistance regulator AmpR, but had no direct effect on expression of the AmpC β-lactamase (Figure 3). We introduced these mutations in the genome of *P. aeruginosa* PAO1 through allelic replacement (Materials and Methods). Since a SNP mutation (G7A) was present at the start of *ampC* gene in one of the alleles, we also made an allelic replacement of this mutation alone in the PAO1 genome to separate the effects caused by the intergenic mutations (Figure S1). For each strain and their isogenic wild type, we measured the Minimal Inhibitory Concentration (MIC) of various β-lactam antibiotics such as imipenem, ceftazidime and ampicillin from carbapenem, cephalosporin and penicillin classes of β-lactams, respectively. For both intergenic alleles, we observed a small but significant increase in the MIC of imipenem and ampicillin (Student *t* test, *P* < 0.01, Figure 4), but not ceftazidime. AmpR regulates β-lactam resistance both through direct activation of AmpC expression as well as via an AmpC-independent manner (26). Irrespective of the mechanism, our results show that acquisition of intergenic mutations between *ampR* and *ampC* is directly linked to a host-relevant phenotypic alteration (*i.e*. reduced β-lactam susceptibility).

**Figure 4:**
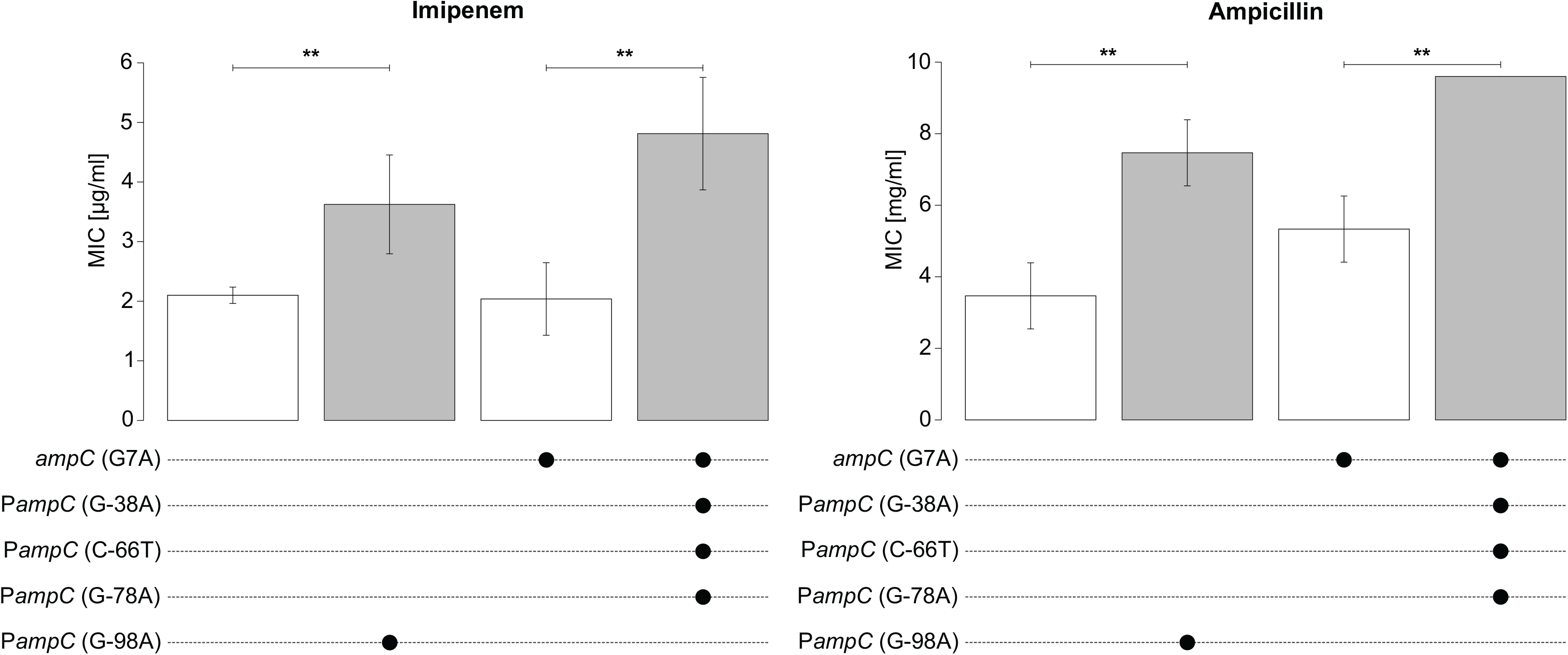
Mutations in the intergenic region between *ampC* and *ampR* cause an increased tolerance towards imipenem and ampicillin. The values for Minimal Inhibitory Concentration (MIC) and the constructed mutations in each strain of PAO1 are shown. Mutation G-98A upstream *ampC* derives from isolate DK2-CF173-1995. Three mutations G-38A, C-66T and G-78A upstream of *ampC* originate from isolate DK1-P43-M2-2002. A SNP mutation at the start of *ampC* (G7A) in DK1-P43-M2-2002 was also constructed in laboratory strain PAO1 to isolate the effect of this mutation and the effect of intergenic mutations from DK1-P43-M2-2002. Error bars indicate standard deviation from three different biological replicates. Double asterisk indicate significant difference between mean MIC of the strains (Two-tailed Student *t* test, *P* < 0.01).

### Intergenic evolution targets essential genes and contributes more to host adaptation than intragenic evolution

Finally, we compared the relative contribution of coding and intergenic mutations to pathogen adaptation. We focused on the large fraction of isolates (n=474) included in this study, in which 52 coding regions were found to be under positive selection during host adaptation (17). In these isolates, we identified 35 pathoadaptive intergenic regions (Materials and Methods, Table S8). Although coding region mutations are numerically dominant over intergenic mutations, normalization to the mutational targets available for intergenic and coding region mutations (89.8% of the *P. aeruginosa* genome contains coding regions) reveals that intergenic regions are 3.7 times more likely to be selected than cording regions and thus appear to play a quantitatively more prominent role in host adaptation (Figure 5a).

**Figure 5:**
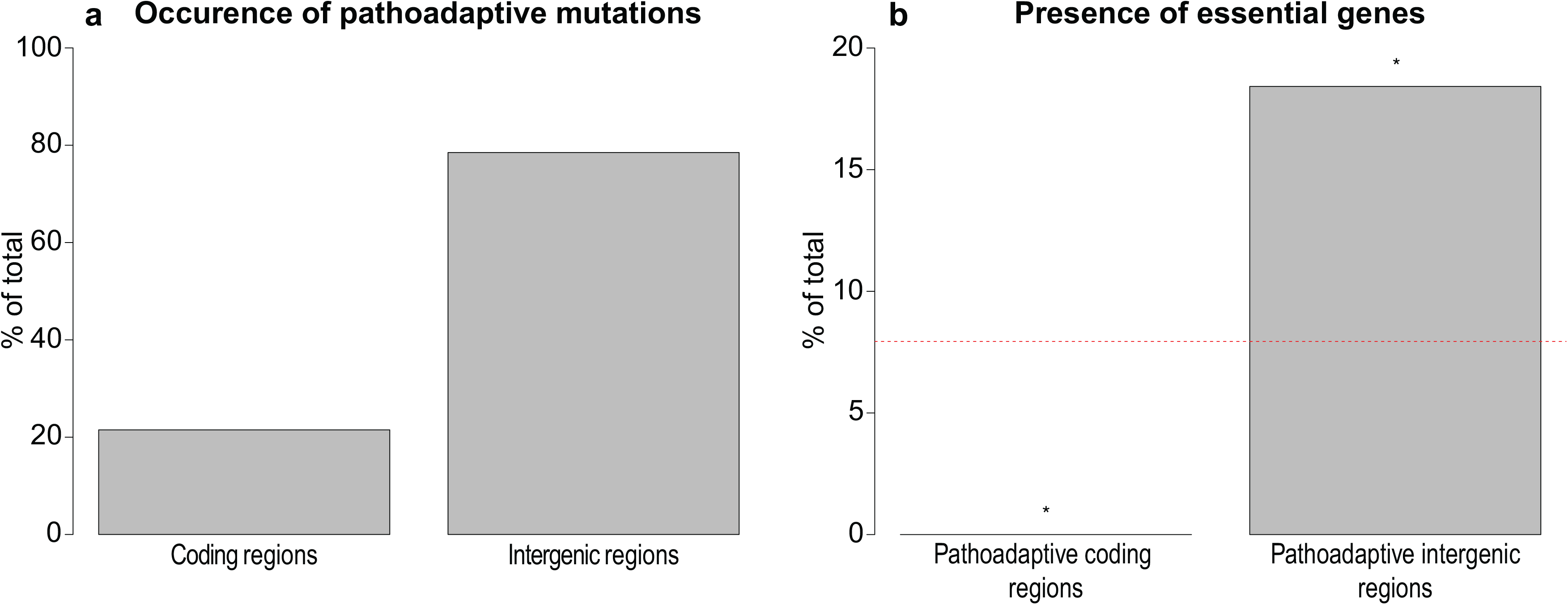
Contribution and function of coding and intergenic mutations. 52 coding regions are under positive selection in a selected subset of our dataset comprising of 474 long-term CF adapted isolates of *P. aeruginosa* (17). In the same isolates, we identify 35 intergenic regions under positive selection for adaptive mutations (Table S8). a) To calculate the relative presence of adaptive mutations within coding or intergenic regions, we divided the number of pathoadaptive features by mutational targets available within each category (89.8% coding and 10.2% intergenic). Final relative frequency for selection of each pathoadaptive feature is shown in the barplot. b) 445 genes are essential for survival of *P. aeruginosa* in CF sputum environment (27). The percentage of these essential genes within 52 pathoadaptive genes and 38 genes downstream of 35 pathoadaptive intergenic regions is demonstrated. Asterisk denotes *P* < 0.05 from two-tailed Fisher’s exact test. Red dashed line indicates the percentage of CF essential genes in PAO1.

We also analyzed qualitative differences between coding and intergenic mutations by determining the presence of essential genes among the pathoadaptive coding and intergenic regions. By cross-referencing the 35 pathoadaptive intergenic regions to the list of 445 genes previously shown to be essential for survival of *P. aeruginosa* PAO1 in CF sputum environment (27), we found that 7 of the 38 genes located immediately downstream of the 35 pathoadaptive intergenic regions are essential (Table S8). Two of these genes (*pyrC* and PA5492) showed altered expression as a consequence of pathoadaptive mutations in their intergenic region, demonstrating that such mutations can indeed modulate expression of essential genes (Figure 3). Importantly, the association between pathoadaptive intergenic regions and essential genes at a level of 18% represents a significant overrepresentation from the normal prevalence of CF sputum essential genes in the *P. aeruginosa* PAO1 genome (Figure 5b, Fisher’s exact test, *P* = 0.029). In contrast, there were no CF sputum essential genes within 52 pathoadaptive coding regions demonstrating a significant underrepresentation of these genes within adaptive coding regions (Figure 5b, Fisher’s exact test, *P* = 0.033).

## Discussion

In this study, we present evidence that intergenic mutations constitute an important part of the genetic basis for host adaptation in *P. aeruginosa*. Generally, the contribution of intergenic regions to evolution of host-adapted variants has received little attention. However, since the development of predictive models of pathogen evolution and discovery of new therapeutic targets (28, 29) rely on understanding the evolutionary response of pathogens to the host environment, identification of the full range of adaptive mutations in both coding and non-coding regions is important. Here, our genome-wide identification of intergenic regions under selection within the host was made possible by combining analysis of parallel evolution across a large number of infected individuals with functional genomics. This approach may be useful for analysis of intergenic regions in connection to host adaptation in other pathogens or niche adaptation in general.

Our identification of pathoadaptive intergenic regions provides insight into the cellular functions targeted by intergenic mutations (Figure 1) and thus points to the selective pressures that confront the pathogen within its CF host. For example, adaptive mutations were found to alter expression of genes such as *cerN* (involved in sphingolipid utilization) (21), *phuR-phuSTUVW* (involved in iron acquisition) (10) and PA4837-34 (involved in zinc acquisition) (23), which strongly indicates that metabolic adaptation for a better exploitation of available nutrients in the host is an important evolutionary driver. Similarly, we observed that mechanisms of the development of tolerance to antibiotics and other inhibitors in the host are also frequent targets of intergenic molecular evolution. Similar functional categories have been found in studies focusing on pathoadaptive mutations within coding regions (14, 16, 17), suggesting that key selective pressures such as nutrient availability and antibiotic stress can be mitigated both by intergenic and coding region evolution in *P. aeruginosa*.

At the functional level, we show that intergenic evolution predominantly targets transcriptional processes to alter the transcriptional activity of downstream genes. However, we also found evidence of parallel evolution in two intergenic small RNAs, four transcriptional terminators, and several cases of mutations located downstream of transcriptional start sites (Table S5 and Table S6), which suggests that adaptive mutations may also target elements that control protein expression at the post-transcriptional level. Importantly, we have shown here and in a previous study that intergenic mutations can be directly responsible for the evolution of important pathogenic traits such as reduced sensitivity to antibiotics (Figure 4) and increased iron uptake (10). Further studies, in particular of pathoadaptive regions upstream of genes with unknown functions, will most likely uncover new mechanisms central to CF host colonization and pathogenesis, and assist in identifying the full complement of stressors present in the host, most of which are currently unknown.

Our study also reveals important qualitative differences between the intergenic and coding region mutations. A generally accepted model is that intergenic mutations would typically confer local and subtle regulatory effects primarily on the immediate downstream genes, whereas mutations in coding regions - with their potential to inactivate entire pathways – would be more likely to cause systemic changes of the physiology of the cell (8, 30). One prediction from this model is that intergenic mutations are associated with less antagonistic effects relative to coding region mutations. While this prediction is difficult to test, our observation of enrichment of essential genes for which intergenic evolution occurred is a clear illustration of this point (Figure 5). The finding that intergenic mutations can bypass the deleterious effects of coding region mutations, thus allowing essential genes to become targets for evolutionary changes, reveals an important aspect of the role of intergenic mutations as well as a key functional difference between intergenic and coding region mutations.

From a quantitative perspective, we find that intergenic mutations are more likely to be selected than coding region mutations during CF host adaptation in *P. aeruginosa* (Figure 5). We hypothesize that the relative contribution of coding and intergenic mutations is variable and depends on a set of identifiable factors of either environmental nature (e.g. niche complexity) or intrinsic to the bacterial pathogen (e.g. genome size, and the number of transcriptional regulatory systems and essential genes encoded in the genome). Although the precise factors that influence the relative contribution of the two types of mutations may be difficult to disentangle, we speculate that in the case of *P. aeruginosa* CF infections, a major contributing factor is the composition of the adaptive environment in the host. The CF host niche is characterized by a complex combination of multiple stressors that must be mitigated for successful bacterial colonization (31). In such environments, mutations in intergenic regions that tune expression levels while maintaining responsiveness to environmental and host-derived cues may result in less pleiotropic effects than mutations that change protein structure or function (30). Further studies of *P. aeruginosa* adaptation in other infections and host environments such as chronic wounds and ulcerative keratitis (32) are required to identify factors that may influence the relative contribution of intergenic and coding region evolution.

Our documentation of the evolutionary significance of intergenic mutations was obtained in the particular genetic and ecological context of *P. aeruginosa* adaptation to the cystic fibrosis airway niche. Nevertheless, our systematic study provides insight into the contribution and functionality of intergenic versus intragenic mutations, which is of broader relevance in connection to bacterial evolution in natural environments. This is supported by recent observations indicating that other contexts may also promote intergenic evolution. For example, our study resonate well with results showing that adaptive intergenic mutations contribute to innovation of novel metabolic functions in laboratory-evolving *E. coli* (33), evidence of a signal of positive selection in *M. tuberculosis* intergenic regions (6) and the suggestion that intergenic evolution may mitigate detrimental fitness effects associated with acquisition of novel genetic material (34). We suggest that adaptive mutations in intergenic regions represent an important but underappreciated aspect of bacterial evolution not only in connection to host colonization but also niche adaptation in other natural environments.

## Materials and Methods

### Bacterial strains and growth conditions

Luria-Bertani (LB) and ABT minimal medium supplemented with 1% glucose and 1% casamino acids (ABTGC) (10) were routinely used for growth of *E. coli* and *P. aeruginosa* strains. *E. coli* CC118 λpir was used for maintenance of recombinant plasmids supplemented with 8 μg/ml of tetracycline. *P. aeruginosa* PAO1 strain was used for phenotypic investigation of intergenic mutations. For final marker selection of *P. aeruginosa*, 50 μg/ml of tetracycline was used.

### Identification of pathoadaptive intergenic regions

The dataset used for analysis of intergenic evolution is described in SI Materials and Methods. Presence of independent clonal lineages of *P. aeruginosa* was verified by recording of MLST allele profiles of each isolate as described in SI Materials and Methods. Pathoadaptive intergenic regions selected across 44 clones were defined as regions containing multiple mutations from isolates of independent clones within a narrow region of less than 30 bp. Similarly, intergenic regions containing multiple distinct mutations from isolates of the same clone within a narrow 30 bp window were also defined as pathoadaptive within that clone. Mutations within narrow windows of pathoadaptive regions were more enriched than what is expected by chance within the dataset. Detailed description is available in SI Materials and Methods. Identification of putative intergenic elements within pathoadaptive intergenic regions is described in SI Materials and Methods.

### Genetic techniques

Genetic engineering of reporter fusion and allelic replacement plasmids are described in SI Materials and Methods. Presence of mutated alleles within reporter fusion and allelic replacement constructs was verified by Sanger sequencing. Reporter fusion plasmids were electroporated into *P. aeruginosa* PAO1 as previously described (35). Constructs with allelic replacements were introduced into *P. aeruginosa* by triparental mating using the helper strain *E. coli* HB101/pRK600 (36). The presence of mutated alleles in tetracycline-sensitive, sucrose-resistant colony isolates was verified by PCR and Sanger sequencing.

### Phenotype assays

Phenotypic expression of reporter fusion strains was investigated as described in SI Materials and Methods. Continuous measurements of growth (OD_600_) and luminescence were recorded by Cytation 5 multimode reader (BioTek) at 200 rpm shaking condition and 37 °C temperature. Lux expression normalized by cell density for all strains was recorded and compared against reporter fusions containing wild type alleles. Measurements were repeated three times for each reporter fusion strain. Determination of MIC values for ceftazidime, imipenem and ampicillin is described in SI Materials and Methods. MIC values were either measured by standard microdilution method in Mueller-Hinton (MH) broth or using E-test on MH agar plates. Measurements were repeated five times for each strain. Statistical differences between means of replicates for reporter fusion and MIC resistance assays were calculated by two-tailed Student *t* test.

## Acknowledgements

We thank Lea M. Sommer and Anders Norman for technical advices in bioinformatics approaches, Esben V. Nisted for help in design of primers, Nicoline Uglebjerg and Caroline A. S. Lauridsen for assistance in identification of putative intergenic elements. We also thank Grith Hermansen, Pavelas Sazinas, Geoff Winsor, Ed Feil, Dominique Schneider and Thomas Hindré for valuable discussions. This work was supported by the Danish Council for Independent Research (6108-00300A) and the Villum Foundation (VKR023113).

## Author contributions

S.M.H.K and L.J. conceived study and designed research. S.M.H.K. performed research. S.M.H.K and L.J. analyzed data and wrote the manuscript.

## References

1. Didelot X, Walker AS, Peto TE, Crook DW, & Wilson DJ (2016) Within-host evolution of bacterial pathogens. Nat Rev Microbiol 14(3):150–162.

2. Hoffman LR, et al. (2009) Pseudomonas aeruginosa lasR mutants are associated with cystic fibrosis lung disease progression. J Cyst Fibros 8(1):66–70.

3. Das S, et al. (2016) Natural mutations in a Staphylococcus aureus virulence regulator attenuate cytotoxicity but permit bacteremia and abscess formation. Proc Natl Acad Sci U S A 113(22):E3101–3110.

4. Honsa ES, et al. (2017) RelA Mutant Enterococcus faecium with Multiantibiotic Tolerance Arising in an Immunocompromised Host. MBio 8(1).

5. Molina N & van Nimwegen E (2008) Universal patterns of purifying selection at noncoding positions in bacteria. Genome Res 18(1):148–160.

6. Thorpe HA, Bayliss SC, Hurst LD, & Feil EJ (2017) Comparative Analyses of Selection Operating on Nontranslated Intergenic Regions of Diverse Bacterial Species. Genetics 206(1):363–376.

7. Kim D, et al. (2012) Comparative analysis of regulatory elements between Escherichia coli and Klebsiella pneumoniae by genome-wide transcription start site profiling. PLoS Genet 8(8):e1002867.

8. Wray GA (2007) The evolutionary significance of cis-regulatory mutations. Nat Rev Genet 8(3):206–216.

9. Osborne SE, et al. (2009) Pathogenic adaptation of intracellular bacteria by rewiring a cis-regulatory input function. Proc Natl Acad Sci U S A 106(10):3982–3987.

10. Marvig RL, et al. (2014) Within-host evolution of Pseudomonas aeruginosa reveals adaptation toward iron acquisition from hemoglobin. MBio 5(3):e00966–00914.

11. Cui Z, et al. (2014) Mutations in the embC-embA intergenic region contribute to Mycobacterium tuberculosis resistance to ethambutol. Antimicrob Agents Chemother 58(11): 6837–6843.

12. Chung JC, et al. (2012) Genomic variation among contemporary Pseudomonas aeruginosa isolates from chronically infected cystic fibrosis patients. J Bacteriol 194(18):4857–4866.

13. Cramer N, et al. (2011) Microevolution of the major common Pseudomonas aeruginosa clones C and PA14 in cystic fibrosis lungs. Environ Microbiol 13(7):1690–1704.

14. Jeukens J, et al. (2014) Comparative genomics of isolates of a Pseudomonas aeruginosa epidemic strain associated with chronic lung infections of cystic fibrosis patients. PLoS One 9(2):e87611.

15. Marvig RL, et al. (2013) Draft Genome Sequences of Pseudomonas aeruginosa B3 Strains Isolated from a Cystic Fibrosis Patient Undergoing Antibiotic Chemotherapy. Genome Announc 1(5).

16. Marvig RL, Johansen HK, Molin S, & Jelsbak L (2013) Genome analysis of a transmissible lineage of pseudomonas aeruginosa reveals pathoadaptive mutations and distinct evolutionary paths of hypermutators. PLoS Genet 9(9):e1003741.

17. Marvig RL, Sommer LM, Molin S, & Johansen HK (2015) Convergent evolution and adaptation of Pseudomonas aeruginosa within patients with cystic fibrosis. Nat Genet 47(1):57–64.

18. Smith EE, et al. (2006) Genetic adaptation by Pseudomonas aeruginosa to the airways of cystic fibrosis patients. Proc Natl Acad Sci U S A 103(22):8487–8492.

19. Stewart AJ, Hannenhalli S, & Plotkin JB (2012) Why transcription factor binding sites are ten nucleotides long. Genetics 192(3):973–985.

20. Stover CK, et al. (2000) Complete genome sequence of Pseudomonas aeruginosa PAO1, an opportunistic pathogen. Nature 406(6799):959–964.

21. LaBauve AE & Wargo MJ (2014) Detection of host-derived sphingosine by Pseudomonas aeruginosa is important for survival in the murine lung. PLoS Pathog 10(1):e1003889.

22. Dasgupta N, Lykken GL, Wolfgang MC, & Yahr TL (2004) A novel anti-anti-activator mechanism regulates expression of the Pseudomonas aeruginosa type III secretion system. Mol Microbiol 53(1):297–308.

23. Mastropasqua MC, et al. (2017) Growth of Pseudomonas aeruginosa in zinc poor environments is promoted by a nicotianamine-related metallophore. Mol Microbiol 106(4):543–561.

24. Gi M, et al. (2015) A novel siderophore system is essential for the growth of Pseudomonas aeruginosa in airway mucus. Sci Rep 5:14644.

25. Toh SM & Mankin AS (2008) An indigenous posttranscriptional modification in the ribosomal peptidyl transferase center confers resistance to an array of protein synthesis inhibitors. J Mol Biol 380(4):593–597.

26. Kumari H, Balasubramanian D, Zincke D, & Mathee K (2014) Role of Pseudomonas aeruginosa AmpR on beta-lactam and non-beta-lactam transient cross-resistance upon pre-exposure to subinhibitory concentrations of antibiotics. J Med Microbiol 63(Pt 4):544–555.

27. Turner KH, Wessel AK, Palmer GC, Murray JL, & Whiteley M (2015) Essential genome of Pseudomonas aeruginosa in cystic fibrosis sputum. Proc Natl Acad Sci U S A 112(13):4110–4115.

28. Yen P & Papin JA (2017) History of antibiotic adaptation influences microbial evolutionary dynamics during subsequent treatment. PLoS Biol 15(8):e2001586.

29. Smith PA & Romesberg FE (2007) Combating bacteria and drug resistance by inhibiting mechanisms of persistence and adaptation. Nat Chem Biol 3(9):549–556.

30. Coombes BK (2013) Regulatory evolution at the host-pathogen interface. Can J Microbiol 59(6):365–367.

31. Folkesson A, et al. (2012) Adaptation of Pseudomonas aeruginosa to the cystic fibrosis airway: an evolutionary perspective. Nat Rev Microbiol 10(12):841–851.

32. Winstanley C, et al. (2005) Genotypic and phenotypic characteristics of Pseudomonas aeruginosa isolates associated with ulcerative keratitis. J Med Microbiol 54(Pt 6):519–526.

33. Blank D, Wolf L, Ackermann M, & Silander OK (2014) The predictability of molecular evolution during functional innovation. Proc Natl Acad Sci U S A 111(8):3044–3049.

34. McNally A, et al. (2016) Combined Analysis of Variation in Core, Accessory and Regulatory Genome Regions Provides a Super-Resolution View into the Evolution of Bacterial Populations. PLoS Genet 12(9):e1006280.

35. Choi KH & Schweizer HP (2006) mini-Tn7 insertion in bacteria with single attTn7 sites: example Pseudomonas aeruginosa. Nat Protoc 1(1):153–161.

36. Kessler B, de Lorenzo V, & Timmis KN (1992) A general system to integrate lacZ fusions into the chromosomes of gram-negative eubacteria: regulation of the Pm promoter of the TOL plasmid studied with all controlling elements in monocopy. Mol Gen Genet 233(1-2):293–301.

